# A mathematical model clarifies the ABC Score formula used in enhancer-gene prediction

**DOI:** 10.1101/2024.11.29.626072

**Authors:** Joseph Nasser, Kee-Myoung Nam, Jeremy Gunawardena

## Abstract

Enhancers are discrete DNA elements that regulate the expression of eukaryotic genes. They are important not only for their regulatory function, but also as loci that are frequently associated with disease traits. Despite their significance, our conceptual understanding of how enhancers work remains limited. CRISPR-interference methods have recently provided the means to systematically screen for enhancers in cell culture, from which a formula for predicting whether an enhancer regulates a gene, the Activity-by-Contact (ABC) Score, has emerged and has been widely adopted. While useful as a binary classifier, it is less effective at predicting the quantitative effect of an enhancer on gene expression. It is also unclear how the algebraic form of the ABC Score arises from the underlying molecular mechanisms and what assumptions are needed for it to hold. Here, we use the graph-theoretic linear framework, previously introduced to analyze gene regulation, to formulate the *default model*, a mathematical model of how multiple enhancers independently regulate a gene. We show that the algebraic form of the ABC Score arises from this model. However, the default model assumptions also imply that enhancers act additively on steady-state gene expression. This is known to be false for certain genes and we show how modifying the assumptions can accommodate this discrepancy. Overall, our approach lays a rigorous, biophysical foundation for future studies of enhancer-gene regulation.

## Introduction

Much of our current understanding of how genes are regulated arose from classical studies in bacteria of the lac operon and *λ*-phage [1]. However, the eukaryotic context differs from the bacterial in many significant ways. One key difference is that, in bacteria, regulatory DNA is found proximal to the gene, typically within 1kb upstream of the transcription start site (TSS), whereas eukaryotic regulatory sequences are found in discrete pieces that may be proximal to, or distal from, the TSS. The eukaryotic regulatory elements known as enhancers form a particularly important class. Enhancers were originally defined as DNA sequences which could drive the expression of genes in a location and orientation independent manner [2, 3]. Since these initial discoveries, many native enhancers have been identified which play critical roles in a variety of processes, such as embryonic development [4], physiology [5] and evolution [6]. Genetic variation in enhancers has also been shown to mediate risk for complex disease [7], Mendelian disease [8] and cancer [9]. Based on these and many other studies, we know that enhancers can be located over 1Mb from a target gene TSS, an individual enhancer may regulate multiple genes, some genes are regulated by multiple enhancers and the set of enhancers actively regulating a given gene may depend on cellular context. These properties have made it difficult to identify the rules governing enhancer-gene regulation.

Given their importance, much attention has been given to systematically identifying enhancer sequences and the genes they regulate. An important breakthrough has been the development of high-throughput CRISPR interference (CRISPRi) screens, which enable putative enhancer sequences to be perturbed in cell culture and the resulting effect on expression of a target gene to be measured [10–13]. These screens typically measure quantitative effects on gene expression as the proportional change in mean gene expression over a cell population. We call this quantity the *fractional change* and, given its importance in this paper, define it formally as follows: let *ψ*(*g*) denote the wild-type mean expression level of a gene *g*, in whatever units are used to measure it, and let 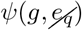 denote the mean expression level of *g* after an enhancer of *g, e*_*q*_, has been perturbed. The *fractional change* of *e*_*q*_ is then the non-dimensional quantity,

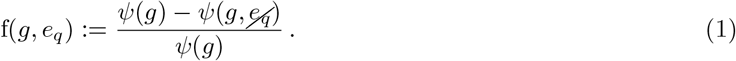

The fractional change for thousands of putative enhancer-gene connections has been measured and computational methods have assessed whether the observed fractional change is statistically different from zero. Current efforts are now focused on two main questions. First, what can we learn about enhancer biology from these screens? Second, can the results from these screens be used to develop computational methods which can predict which enhancers regulate which genes in arbitrary cellular contexts?

The Activity-by-Contact (ABC) model has been proposed as a way to make progress on both of these questions [11]. The ABC model is based on the mechanistic notion that an enhancer’s effect on gene expression depends on the intrinsic strength of the enhancer (activity) and the frequency with which it comes into physical proximity to the gene promoter (contact). The ABC model gives rise to the ABC Score, a quantitative formula which is intended to predict the fractional change observed in an enhancer perturbation experiment. For a gene, *g*, with *N* putative enhancers, *e*_1_, · · ·, *e*_*N*_, the ABC Score for a specific enhancer *e*_*q*_, is given by,

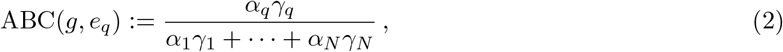

where *α*_*i*_ represents the activity of *e*_*i*_ and *γ*_*i*_ represents the contact frequency between *e*_*i*_ and the promoter of

*g*. In [11] a putative enhancer was defined as a chromatin-accessible DNA element of approximately 500 base pairs; *α*_*i*_ was assigned using measures of chromatin state of the enhancer, such as DNase-Seq and H3K27ac ChIP-Seq; *γ*_*i*_ was assigned using the contact frequency between a putative enhancer and the gene promoter, as measured by Hi-C; and the sum in the denominator of Eqn.2 was taken over all putative enhancers within 5Mb of *g*.

The ABC Score is reasonably effective at predicting the results of CRISPRi screens. When considered as a binary classifier, the ABC Score has achieved a precision of 59% at 70% recall benchmarked against a database of nearly 4,000 putative enhancer-gene connections in the K562 cell line [11]. Similar performance has also been observed in other cell types [11, 14] and in subsequent benchmarking against other CRISPRi screens in K562 cells [15, Fig.S8a]. We emphasize that the ABC Score is computed directly from genomic data orthogonal to the CRISPRi experiment. As such, it has no free parameters and does not require fitting or training. The classification ability of the ABC Score and its modest input data requirements have resulted in its widespread use to interpret non-coding genetic variation [14, 16, 17], identify enhancers in disease related contexts [18, 19] and investigate the dosage effect of transcription factor concentration on gene expression [20].

Despite its practical utility as a binary classifier, the ability of the ABC Score to predict the fractional change is fundamentally limited [11, Fig.3c]. From Eqn. 2, it is clear that the sum of the ABC Scores over all putative enhancers of a given gene is equal to 1,

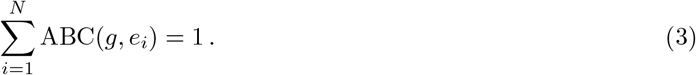

We define the total fractional change of a gene to be the sum of the fractional changes of all enhancers for the gene, f(*g, e*_1_) + · · · + f(*g, e*_*N*_). If the ABC model were perfectly reflecting the fractional change, so that ABC(*g, e*_*i*_) = *f* (*g, e*_*i*_), it would predict that the total fractional change for all genes is equal to 1. However, experimentally, a range of total fractional changes has been observed from 0 to greater than 3 [10, 11, 14, 21–23]. This incompatibility is a consequence of the algebraic structure of the ABC Score formula and cannot be resolved in a straightforward way. For example, it cannot be resolved by using different types of epigenomic data to assign values to *α*_*i*_ or *γ*_*i*_.

What, if anything, about enhancer biology can be concluded from the successes and limitations of the ABC Score? We believe that considering this question requires a formal description of the ABC model. The original description of the ABC model is *informal*, in the sense that the relationships between the mechanisms of activity and contact and the ABC Score formula were not determined by formal mathematical arguments. In consequence, the biological and biophysical assumptions that underlie formulas of this kind have not been clarified.

In the present paper, we present a strategy for the formal mathematical modeling of enhancer-gene regulation. We introduce the *default model*, a set of assumptions for how multiple enhancers independently regulate a gene. We show that a formula with the same algebraic structure as the ABC Score formula in Eqn.2 can be rigorously derived from a special case of the default model. This clarifies the assumptions that underlie the ABC Score formula. However, these assumptions also imply that the total fractional change of a gene is equal to one. We show how changing the assumptions of the default model can lead to total fractional changes which are less than or greater than one. More generally, the framework introduced here offers a rigorous foundation for future studies of enhancer-gene regulation.

## Results

### An activation-communication model of enhancer function

Our approach to modelling enhancer-gene regulation is based on the linear framework, a method of using graphs to analyse biomolecular systems [24–26] that has been previously introduced to study gene regulation [26]; see [27, 28] for up-to-date reviews. The graphs in question have vertices that are linked by labelled, directed edges. The vertices represent molecular states of DNA, the edges represent transitions between these molecular states and the labels represent the transition rates, which are positive numbers with dimensions of (time)^−1^.

An example linear framework graph is shown in Fig.1a. This graph, which we have called *H*, represents a single enhancer which can be either activated (filled red circle) or not and in communication with its target gene (curved arrow) or not. It thereby captures the two main notions in the original ABC model, although we prefer to speak here of “communication” rather than of “contact” (see below). These two features of the enhancer are treated in the graph as being independent of each other: the rates for becoming activated or deactivated do not depend on the state of communication, and the rates for making or losing communication do not depend on the state of activation. Independence will be one of the central features of our treatment and will appear both in how an individual enhancer is treated, as in this example in Fig.1a, and in how a gene is regulated by multiple enhancers, as we will explain below.

**Figure 1:**
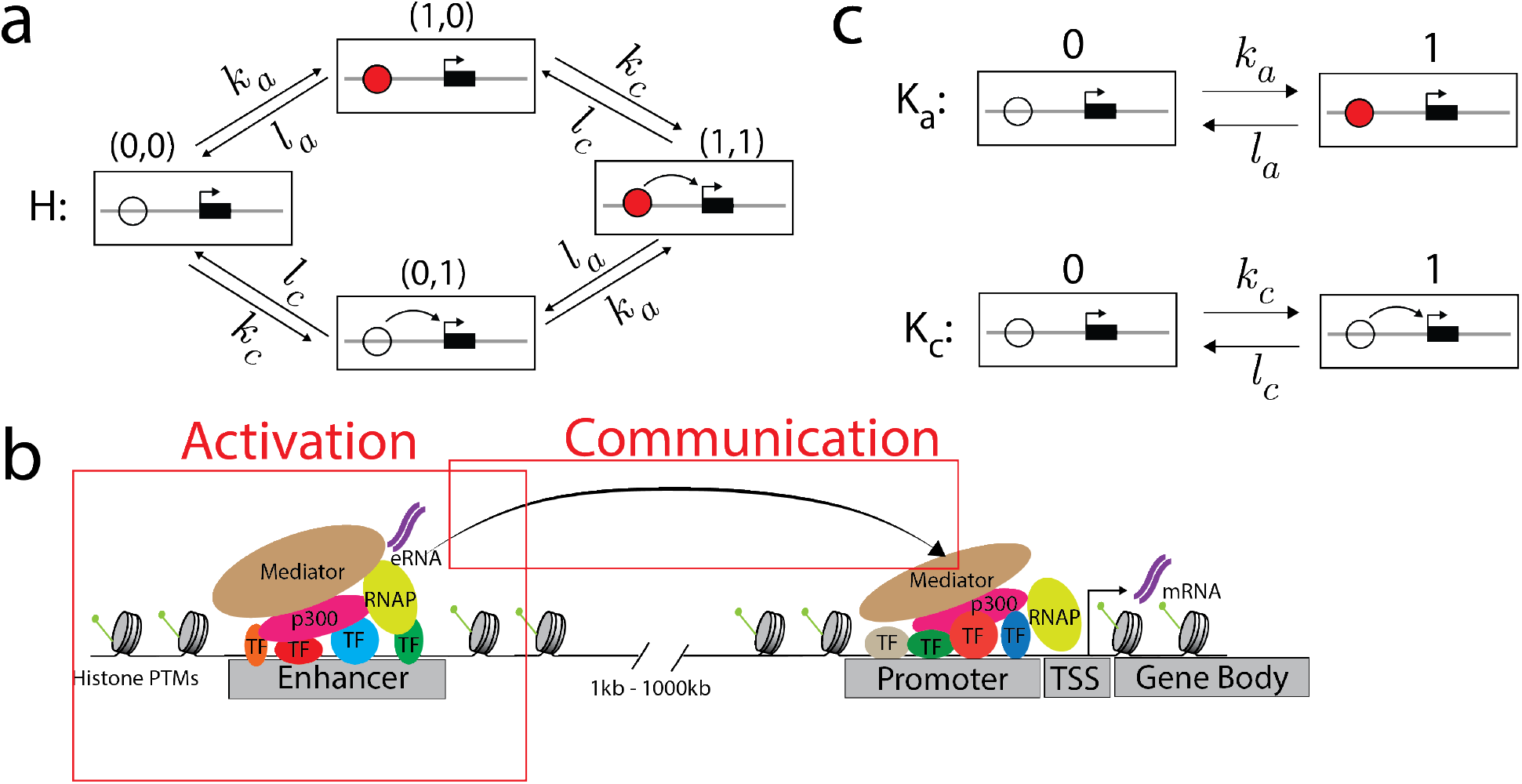
The activation-communication coarse graining. **a)** An example linear framework graph, *H*, representing a coarse-grained view of an enhancer. Each vertex contains a schematic of the enhancer (circle) and its target gene (black rectangle with the transcription start site marked with an arrow). The enhancer may be activated (filled red circle) or communicating (curved arrow to the target gene), encoded in the notation (*i, j*) used to denote vertices. The edge labels show that activation and communication take place independently of each other. **b)** A more detailed picture of the molecular complexity that may underlie the coarse-grained graph in panel **a**, as described further in the text. **c)** The example graph *H* in panel **a** is the graph product of two simpler 2-vertex graphs, *K*_*a*_, which represents activation, and *K*_*c*_, which represents communication. The product structure of *H* is equivalent to the independence of activation and communication.

The graph *H* in Fig.1a represents a *coarse-graining* of the actual complexity of enhancer-gene regulation (Fig.1b). Activation is intended to capture processes local to the enhancer sequence such as transcription factor binding, chromatin reorganisation, nucleosome remodelling, recruitment of co-regulators or transcription of the enhancer sequence itself to generate enhancer RNA. Communication refers to the processes by which information is transferred from the enhancer to its target gene. Many communication mechanisms have been proposed including physical contact through DNA looping [29], diffusion of regulatory molecules and phase separation [31]. We thus use the word ‘communication’ instead of ‘contact’ to reflect that enhancer-gene regulation may not require physical contact. It is, of course, possible that the specific activation or communication mechanisms may differ between enhancers. The value of this coarse-graining lies in not making commitments about the underlying mechanism, at the price of ignoring the potential consequences of how activation and communication are implemented in molecular terms. This particular coarse-graining will facilitate our clarification of the ABC Score formula below.

Having provided an example of a linear framework graph and explained how it describes the biological context that we will be studying, we now go into the details of the linear framework. We will make use of the example in Fig.1a throughout this work.

### Preliminaries on the linear framework

#### Notation and terminology

We will start by introducing some basic ideas about linear framework graphs. We will use a letter like *G* or *H* to refer to a graph. Vertices will generally be denoted *i, j*, etc. We will use the notation *i* ∈ *G* to mean the state *i* from the graph *G*. Edges will be denoted *i* → *j* and edge labels will be denoted *ℓ*(*i* → *j*). So, using the notation for the example graph *H* in Fig.1a, *ℓ*((0, 0) → (0, 1)) = *ℓ*((1, 0) → (1, 1)) = *k*_*c*_ (the notation for the vertices in this graph arises from its product structure and will be explained later). If some feature *X* is being discussed for different graphs, we will sometimes use brackets, as in *X*(*G*), or a subscript, as in *X*_*G*_, to specify the graph in question. We will use the word *structure* to refer to just the vertices and edges of a graph, ignoring the edge labels; when we say “graph”, we will always be including the labels, even when they are not mentioned explicitly.

#### The Markov process

A graph *G* is equivalent to a finite-state, continuous-time, time-homogeneous Markov process [25, 28, 32]. This stochastic behaviour can be understood as follows. If the system is in state *i*, then for each edge *i* → *j* which leaves *i*, a “firing” time is randomly chosen from the exponential probability distribution, *λ* exp(−*λt*), where *λ* is the transition rate of that edge, *λ* = *ℓ*(*i* → *j*), and the edge with the lowest firing time is taken, at that time. This generates a stochastic trajectory of states and transitions. If we follow a trajectory up to time *T* and measure the proportion of time spent in state *i*, then that ratio stabilises with increasing *T* to become the *steady-state probability* of state *i* [32], which we will denote by 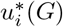. Provided *G* is *strongly connected*, this quantity does not depend on the state in which the trajectory starts and the steady-state probability is a property of the graph [25]. A strongly connected graph is one in which any two distinct vertices, *i* and *j* ≠ *i*, are connected by a directed path, *i* = *i*_1_ → *i*_2_ → · · · → *i*_*k*_ = *j*. The example graph *H* in Fig.1a is strongly connected but ceases to be if the edges (1, 1) → (1, 0) and (1, 1) → (0, 1) are removed. We will assume from now on that all our graphs are strongly connected.

#### Thermodynamic equilibrium and steady-state probabilities

One of the advantages of the linear framework is that, provided the graph is finite, its steady-state probabilities can be calculated algebraically in terms of the edge labels. (We will encounter an infinite graph below but, as we will see, we do not have to deal with them directly and can work only with finite graphs.) If the graph can reach *thermodynamic equilibrium* the algebra can be done quite easily but, importantly, it can also be done when the graph is away from thermodynamic equilibrium, although the formulas become more complicated. A graph can reach thermodynamic equilibrium if, and only if, it satisfies two conditions. First, it must be *reversible*, so that if there is an edge *i* → *j*, then there is also an edge *j* → *i*, which represents the reverse of the process that corresponds to *i* → *j*. Second, it must satisfy the *cycle condition*: the product of the label ratios around any cycle of reversible edges must be 1. The graph in Fig.1a is evidently reversible and has only one cycle of reversible edges, (0, 0) ⇋ (1, 0) ⇋ (1, 1) ⇋ (0, 1) ⇋ (0, 0), for which the product of label ratios is

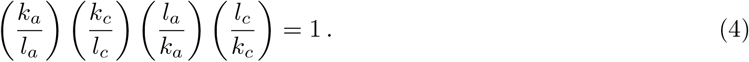

For this graph, the independence of activation and communication ensures that the graph can reach thermodynamic equilibrium.

When a graph can reach thermodynamic equilibrium, its steady-state probabilities can be calculated as follows. First, choose any vertex as a reference; let us call it 1. Second, choose any path of reversible edges from 1 to the state in question, say *i*: 1 ⇋ *i*_1_ ⇋ · · · ⇋ *i*_*k*_ = *i*. The steady-state probability of *i* is then proportional to the product of the label ratios along this path,

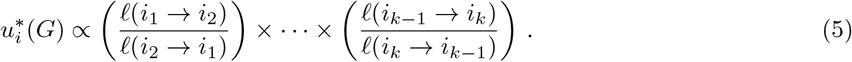

It is a simple consequence of the cycle condition that the quantity on the right-hand side of Eqn.5 does not depend on the choice of path from 1 to *i*. The proportionality constant in Eqn.5 is readily obtained by exploiting the fact that the sum of all the probabilities must be 1, so that, if the vertices are denoted 1, · · ·, *N*, then 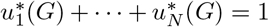 If we follow this prescription for the graph in Fig.1a, we find that, for example,

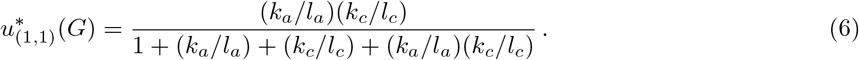

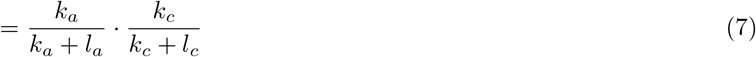

(We will sometimes use a “·” to denote multiplication to make formulas like this look clearer.) The reorganisation of Eqn.6 into Eqn.7 reveals a product structure in the algebra whose significance will emerge below. The formula in Eqn.6 is the same as would arise from equilibrium statistical mechanics. It is one of the features of the linear framework that it reduces to equilibrium statistical mechanics for systems that are at thermodynamic equilibrium but also yields algebraic formulas for systems away from thermodynamic equilibrium.

#### Product graphs as models of independence

In studying gene regulation, a very helpful construction is that of a *product graph*, because it captures the default situation in which two or more genetic systems operate independently of each other. The example graph in Fig.1a is a case in point. This graph *H* is the product of the graphs *K*_*a*_ and *K*_*c*_ in Fig.1c. Here, *K*_*a*_ is a two-vertex graph that represents just the activation of the enhancer and *K*_*c*_ is a two-vertex graph that represents just the communication.

We will use *K*_*a*_ and *K*_*c*_ to describe the product graph construction. We will do this in two steps. We will first specify the vertices and edges by building the *product structure*, denoted *K*_*a*_ *× K*_*c*_, and then we will specify the labels to get the *product graph*, denoted *K*_*a*_⊗*K*_*c*_. As we will see below, product structures underlie other constructions in which the independence of the product graph is broken, which is why it is helpful to distinguish structures and graphs. The vertices in *K*_*a*_ *× K*_*c*_ are ordered pairs, (i, j), of vertices *i* ∈ *K*_*a*_ and *j* ∈ *K*_*c*_. The edges in *K*_*a*_ *× K*_*c*_ arise from the edges in either component *K*_*a*_ or *K*_*c*_, taken independently of the state of the other component. In other words, if *i*_1_ → *i*_2_ is any edge in *K*_*a*_, then (*i*_1_, *j*) → (*i*_2_, *j*) is an edge in *K*_*a*_ *× K*_*c*_, for all *j* ∈ *K*_*c*_; similarly, if *j*_1_ → *j*_2_ is any edge in *K*_*c*_, then (*i, j*_1_) → (*i, j*_2_) is an edge in *K*_*a*_ *× K*_*c*_, for all *i* ∈ *K*_*a*_; these are the only edges in *K*_*a*_ *× K*_*c*_. This prescription yields the structure of the graph *H* in Fig.1a.

The labels of the product graph, *K*_*a*_ ⊗ *K*_*c*_, are also inherited from those in *K*_*a*_ or *K*_*c*_, independently of the state of the other component,

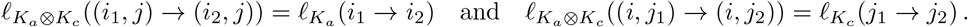

We see that *K*_*a*_ ⊗ *K*_*c*_ corresponds exactly to the graph *H* in Fig.1a. The graph product precisely captures the sense in which the components of the product, here *K*_*a*_ and *K*_*c*_, operate independently of each other: the transitions in either component are unaffected, as to their occurrence and their rates, by the state of the other component.

In the more general case of a product of *m* graphs, the vertices are naturally indexed as ordered tuples, (*i*_1_, · · ·, *i*_*m*_).

One of the consequences of the product graph construction is that its steady-state probabilities are easily calculated. If *K*_1_, · · ·, *K*_*N*_ are any set of *N* strongly connected graphs, then the steady-state probabilities in the product graph *K*_1_ ⊗ · · · ⊗ *K*_*N*_ can be computed by multiplying the steady-state probabilities in the individual graphs,

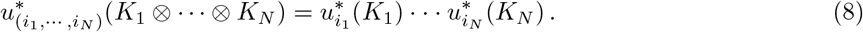

Eqn.8, which is proved in [26], again captures the sense in which the components *K*_1_, · · ·, *K*_*N*_ are independent of each other. We note that Eqn.8 holds even for graphs which are unable to reach thermodynamic equilibrium.

We can see Eqn.8 at work for the graph in Fig.1a, which is the product of the graphs *K*_*a*_ and *K*_*c*_ in Fig.1c. If we follow the prescription in Eqn.5, we see that

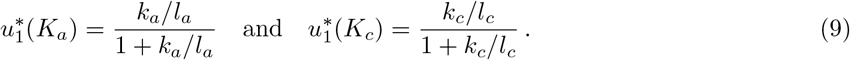

If we apply Eqn.8 to the formulas above, we see that,

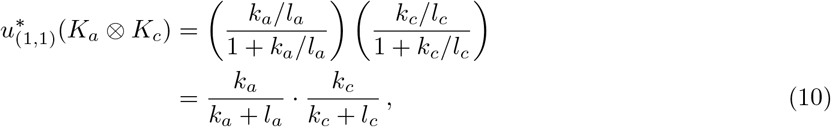

which recovers the expression in Eqn.7, whose algebraic product structure is now seen to reflect the underlying product graph.

Eqn.8 for individual vertices has a straightforward extension to subsets of vertices. To explain this, let *K* be any graph and let *S* ⊆ *K* be any subset of vertices in *K*. The steady-state probability of being in any vertex of *S*, denoted 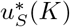. is given by 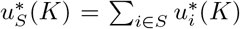. Now suppose, as above, that *K*_1_, · · ·, *K*_*N*_ are any strongly connected graphs. Let *S*_*i*_ ⊆ *K*_*i*_ be any subset of vertices of *K*_*i*_ and let *S*_1_ *×* · · · *× S*_*N*_ be the corresponding *set product* in *K*. This set product, for which we use, for convenience, the same notation as for the product structure, has the obvious definition that it consists of all those tuples (*i*_1_, · · ·, *i*_*N*_) where *i*_*k*_ ∈ *S*_*k*_. It is then a simple consequence of Eqn.8 that,

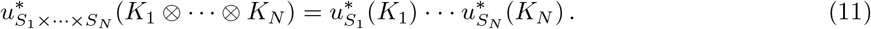

One of the implications of Eqn.11 is that if we take *i*_*j*_ to be a coordinate that runs over the vertices of *K*_*j*_, then the probability that *i*_*j*_ has a particular value, say *i*_*j*_ = *b*, remains the same irrespective of the other factors in the graph product,

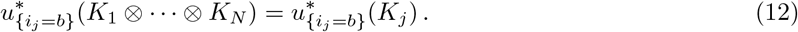

Eqn.12 follows from Eqn.11 because the subset {*i*_*j*_ = *b*} in *K*_1_ ⊗ · · · ⊗ *K*_*N*_ is the product subset,

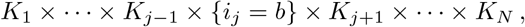

and 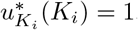. We can see an example of Eqn.12 at work in Figure 1. Let {a=1} = {(1,0), (1,1)} be the subset of vertices of *H* in which the enhancer is activated. Then Eqn.12 shows that 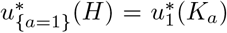. We will make further use of Eqn.12 in what follows.

#### The gene expression response

The graphs we have considered up to now are models of the regulatory state of the gene. We now discuss how to incorporate the production and degradation of mRNA. The standard approach in the literature is known as *kinetic modeling* and uses a Markovian framework based on the chemical master equation [33]. We follow this same approach within the graph-theoretic setting introduced here.

At any given time, the state of gene expression is specified by a certain number of molecules of the corresponding mRNA. This number increases by 1 each time RNA polymerase transcribes the gene and decreases by 1 each time an mRNA molecule is degraded or lost through transport out of the nucleus. We can represent such an expression system by the (semi)-infinite *pipeline* structure, *P*, in which the state *p* represents the number *p* of mRNA molecules, from *p* = 0 onwards, and the edges correspond to mRNA production, *p* → *p* + 1, and degradation or loss, *p* → *p* − 1 (Fig.2a).

**Figure 2:**
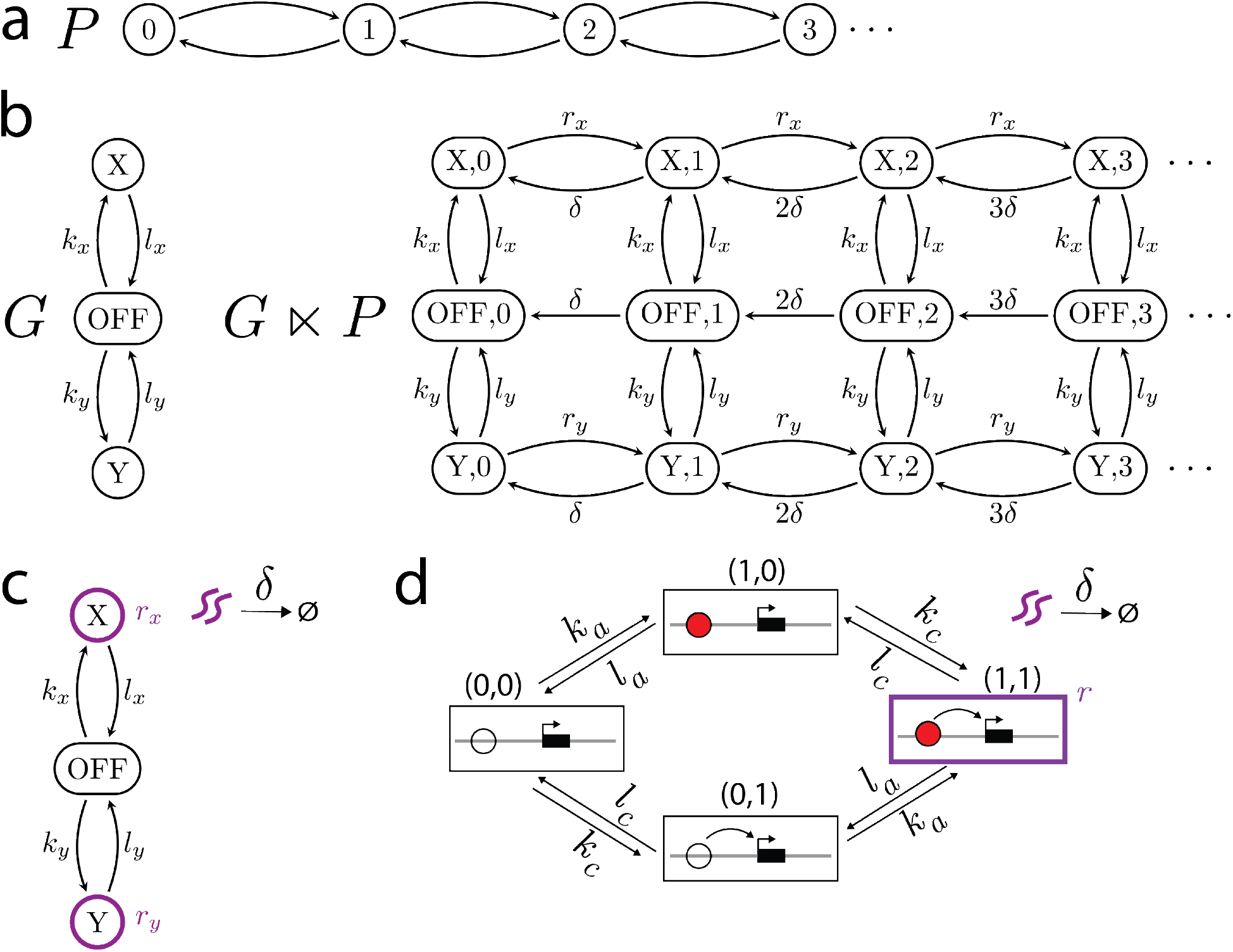
Modeling mRNA production and degradation through copy-number graphs. **a)** The pipeline structure *P* represents the number of mRNA molecules and their production and loss. **b)** An example regulatory graph, *G*, and the resulting copy-number graph *G* ⋉ *P*. In this example *G* has two production states, *X* and *Y*, with corresponding mRNA production rates *r*_*x*_ and *r*_*y*_ respectively. We note that *G* ⋉ *P* is a sub-structure of *G* × *P* ; it has the same vertices but lacks the edges corresponding to a production rate of zero. *G* and *P* also do not operate independently in *G* ⋉ *P* because the mRNA production rates depend on the regulatory state. **c)** A compact way to represent *G* ⋉ *P*. The production states are outlined in purple with corresponding mRNA production rates. The degradation rate, *δ*, is shown above the arrow from purple squiggles (mRNA) to the empty set ∅. **d)** The compact representation of the graph *H* ⋉ *P*, where *H* is given in Fig.1a. The only production state is the state (1, 1), in which the enhancer is both activated and communicating, which has production rate *r*.

Given a gene-regulatory graph, *G*, we represent the overall system of regulation and expression by a *copy-number graph, G* ⋉ *P*, that will be derived from the product structure, *G × P* (Fig.2b). The states of *G* ⋉ *P* are identical to those of *G × P* but *G* ⋉ *P* may not have all the edges that are present in *G × P*. Each state in *G* ⋉ *P* keeps track of the regulatory state of the gene and the number of mRNA molecules that are present. We now discuss how to assign labels to this graph (Fig.2b). We assume that each state, *i* ∈ *G*, has a corresponding non-negative rate of mRNA production, *r*_*i*_(*G*) ≥ 0. If *r*_*k*_(*G*) = 0, so that mRNA production is not possible in state *k* of *G*, then the edges (*k, p*) → (*k, p* + 1) are removed from *G × P* for all *p* ∈ *P*. (Note that edge labels must always be positive.) This is the only way in which the structure of *G* ⋉ *P* differs from that of *G × P*. If *r*_*k*_(*G*) *>* 0, we will assume that the rate of mRNA production does not change with the number of mRNA molecules that have been expressed, so that *ℓ*((*k, p*) → (*k, p* + 1)) = *r*_*k*_(*G*) for any *p* ∈ *P*. As for mRNA degradation or loss, this takes place independently of the regulatory system, so the most parsimonious assumption is that its rate is proportional to the number of mRNAs that are present and is independent of the regulatory state. Accordingly, we may write *ℓ*((*k, p*) → (*k, p* − 1)) = *δ*(*G*) · *p* for any *k* ∈ *G* and any positive *p* ∈ *P*, where *δ*(*G*) is the degradation rate constant. Finally, we assume that regulatory transitions do not depend on gene expression, so that *ℓ*((*i, p*) → (*j, p*)) = *ℓ*_*G*_(*i* → *j*) for all *i, j* ∈ *G* and for all *p* ∈ *P*. A compact way to visually represent a copy-number graph is shown in Fig.2c.

At steady state, *G* ⋉ *P* gives rise to a probability distribution over the mRNA copy number. We will define the response of the gene, which we will denote by *R*(*G*), to be given by the average of this number

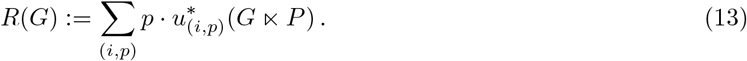

We note that *R*(*G*) ≥ 0.

Because *G* ⋉ *P* is not a finite graph, the prescription given in Eqn.5 for calculating steady-state probabilities no longer works. (*G* ⋉ *P* is also not at thermodynamic equilibrium, unless every regulatory state has the same rate of mRNA production, as can be checked by following the cycle condition formula in Eqn.4.) However, we can appeal to a very useful theorem, due to Sanchez and Kondev, which tells us that we do not have to operate on *G* ⋉ *P* in order to calculate *R*(*G*) [34]. Translating their work into the graph-theory language used here, we find that the response of the gene can be calculated in terms of the average of *r*_*i*_(*G*) over only the the steady-state probabilities of *G*, normalized by *δ*(*G*),

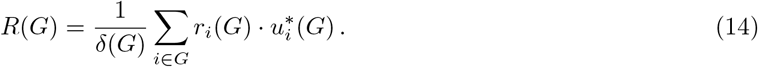

It follows that, although infinite graphs arise to represent the mRNA expression system, we do not need to work with them to calculate the mean steady-state expression *R*(*G*), under the assumptions made above. Sanchez and Kondev did not use graph theory in their work, so we provide an independent graph-based proof of Eqn.14 in the Methods. In subsequent work, we will show how the copy-number graphs introduced here lead to generalizations of the results of [34] but we do not need that for the present paper.

This result provides some justification for reducing the notational clutter from multiple instances of *P*. We will refer to the *regulatory graph G* when we mean *G* on its own, and to the *copy-number graph G* when we mean *G* ⋉*P*, defined for some specified choice of production rates *r*_*i*_(*G*) and degradation rate *δ*(*G*). These parameters may not be explicitly mentioned when speaking of a copy-number graph but they should be kept in mind.

As an illustration of Eqn.14, we will assign production rates to the graph in Fig.1a and compute its response. We will make the assumption that mRNA is only produced when the enhancer is both activated and communicating (Fig. 2d). The mRNA production rates of *H* are therefore given by

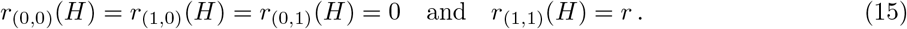

We can now use Eqn.14 to calculate the response, *R*(*H*), taking advantage of Eqn.10, in which we exploited the product graph decomposition *H* = *K*_*a*_ ⊗ *K*_*c*_. We see that,

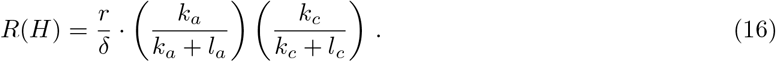

It follows from Eqns.9 and 12 that we can interpret *k*_*a*_*/*(*k*_*a*_ + *l*_*a*_) as the probability that the enhancer is activated and, similarly, *k*_*c*_*/*(*k*_*c*_ + *l*_*c*_) as the probability that the enhancer is communicating. Eqn.16 tells us that the response of *H* is the product of the ratio of production to degradation, the probability of activation and the probability of communication.

This concludes our analysis of a gene regulated by a single enhancer using the activation-communication coarse graining. We now turn to considering how multiple enhancers work together to regulate gene expression.

### A default model of how multiple enhancers independently regulate a gene

We now introduce the *default model* of enhancer-gene regulation. This is a set of assumptions for how multiple enhancers collectively regulate a gene in an independent manner. We previously introduced the product graph construction which represents independence between regulatory graphs. We now broaden those assumptions to also allow for mRNA production. We expect this default model construction to be of general interest. In the next section we will show how a special case of the default model clarifies the ABC Score formula.

Consider a gene, *g*, that is regulated by *N* enhancers, *e*_1_, · · ·, *e*_*N*_. We will assume that enhancer *e*_*l*_ is modelled by the graph *G*_*l*_. We make no assumptions about *G*_*l*_ other than the prevailing assumption that all our graphs are strongly connected. *G*_*l*_ could be substantially more complicated than the graph in Fig.1a and could incorporate, for example, chromatin organisation, nucleosomes, co-regulators, post-translational modifications, chromosome conformation, etc [26]. In particular, there is no requirement that *G*_*l*_ should be able to reach thermodynamic equilibrium. At this point our assumptions are very general and could apply to essentially any enhancer, when considered from a Markovian perspective.

We denote the graph that models the collective regulation of the enhancers by *G* and describe how *G* is defined in terms of the *G*_*l*_.

The first assumption says that each enhancer has its own individual effect.

1. Individuality. Each enhancer *e*_*l*_, when acting in the absence of any of the other enhancers, drives gene expression at the rate *r*_*i*_(*G*_*l*_) ≥ 0 for each state *i* ∈ *G*_*l*_, and gives rise to the response *R*(*G*_*l*_), as defined by Eqn.13. If the enhancer is unable to drive expression on its own, then *r*_*i*_(*G*_*l*_) = 0 for every state *i* ∈ *G*_*l*_. The next two assumptions specify how the enhancers work together.
2. Regulatory independence. Each enhancer acts independently of all the others, so that the regulatory graph of *G* is given by the product graph *G*_1_ ⊗ · · · ⊗ *G*_*N*_.
3. Production-rate summation. Each enhancer independently influences mRNA production. Accordingly, if (*i*_1_, · · ·, *i*_*N*_) is a state in *G*, then its mRNA production rate is a sum of the corresponding production rates in each enhancer graph:

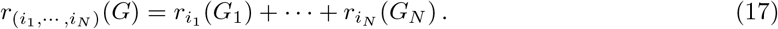 The summation of rates in Assumption 3 arises for the following reason. If each enhancer influences mRNA production independently, then the time at which an mRNA is produced will be the minimum of the times at which each individual enhancer has its effect on production. These individual times are exponentially distributed with rates 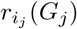 for state *i*_*j*_ in *G*_*j*_. The minimum of several exponentially distributed random variables is a random variable that is also exponentially distributed, with rate given by the sum of the individual rates. This leads to Eqn.17. The final assumption specifies the degradation rates.
4. Uniform degradation. Since mRNA degradation is a separate process to gene regulation and gene expression, we consider the characteristic degradation rate to be a property of the gene, not the enhancer. As such, each graph *G*_*l*_ is assumed to have the same degradation rate, *δ*(*G*_*l*_) = *δ* for all *l*, and the mRNA degradation rate of *G* is also *δ*: *δ*(*G*) = *δ*.

For any set of copy-number graphs *G*_1_, …, *G*_*N*_, we denote the copy-number graph which models their collective effect on transcription according to Assumptions 1 to 4 by the graph

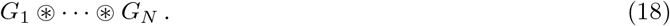

Example constructions using the default model are given in Fig.3.

**Figure 3:**
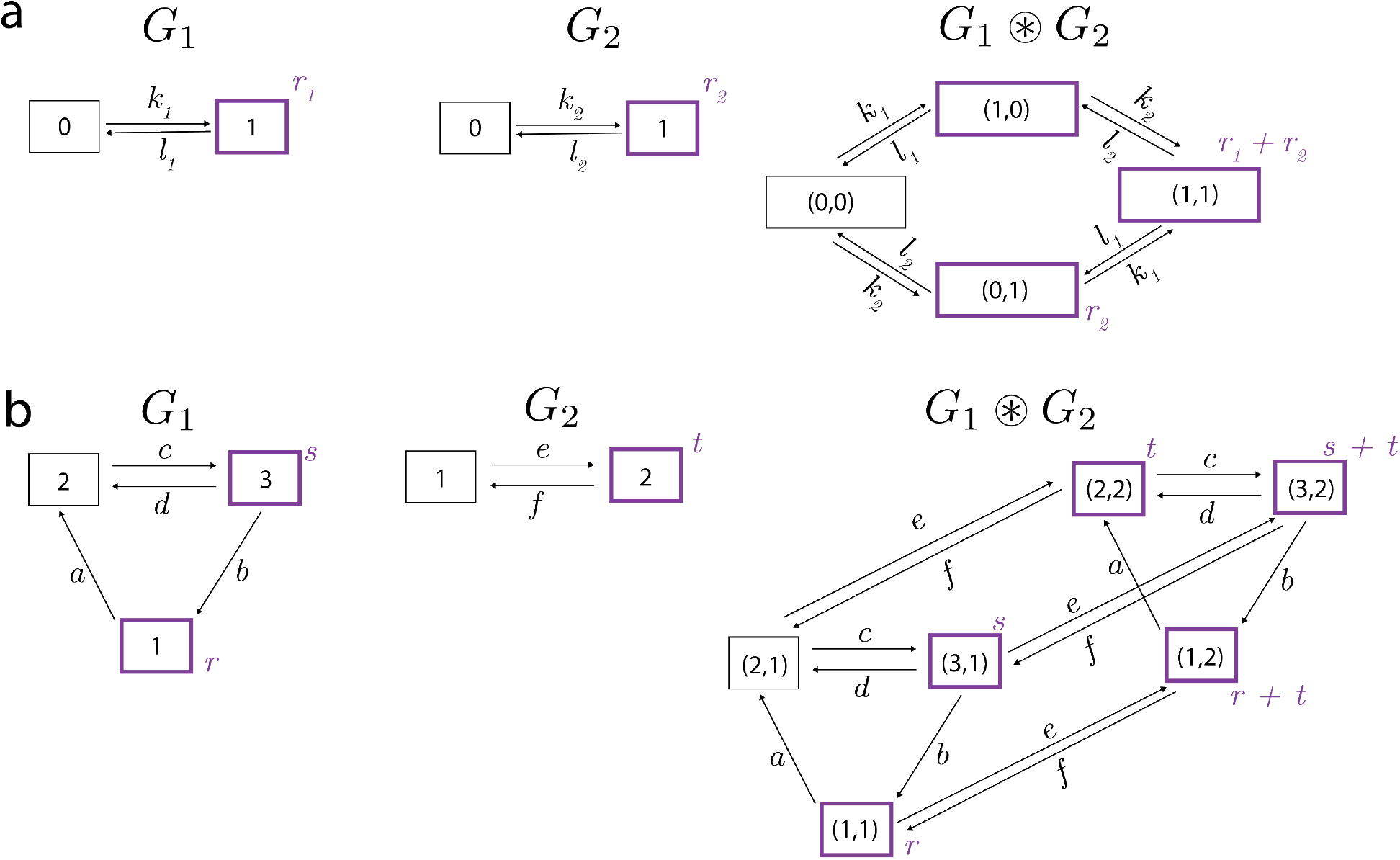
Two examples **a** and **b** of the default model construction. The copy-number graphs are depicted in compact format, as shown in Fig.2c but omitting the degradation symbols for clarity. Production states are outlined in bold purple with corresponding production rates in purple text. The model in **b** is adapted from Figure 9 of [26].

Assumptions 1 to 4 specify our default model of how enhancers collectively regulate a gene. Whether any of the default model assumptions hold for an individual gene is a question that has to be addressed experimentally. In particular, we would expect that Assumption 3 would eventually break down as more enhancers are added to a gene since production rates will be limited by the physical processes involved in transcription. Our goal here is to rigorously work out the consequences of these assumptions, so that we know what to expect when the assumptions do hold and can compare these predictions to what is found experimentally. Of particular significance is that the assumptions above imply that the collective response of the enhancers is always the sum of their individual responses,

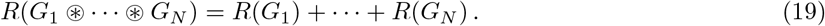

A proof of this fundamental property of the default model is given in the Methods.

A recent commentary has argued that formal definitions and rigorous modeling are necessary to investigate whether a set of enhancers is “greater than the sum of its parts” [35]. We fully agree and suggest that the notion of independence encoded by the default model, which gives rise to Eqn.19, could serve as a definition of what it means for a gene to be the *sum of its parts*.

Transcription in the default model relies on the presence of enhancers. It is well known that the promoter sequences at some eukaryotic genes are sufficient to drive transcription even in the absence of distal enhancers [36, 37]. It is a future area of research to incorporate the role of core promoter elements and promoter proximal regulatory sequences along with their interactions with distal enhancers.

### A clarification of the ABC Score formula

#### Enhancer perturbation and deletion fidelity

To see how formulas similar to the ABC Score can be derived from the default model, we need to consider how to formally model perturbations to enhancers such as genetic deletions or CRISPRi. As previously, we will assume that the target gene *g* is collectively regulated by enhancers *e*_1_, · · ·, *e*_*N*_. We assume that enhancer *e*_*i*_ is modeled by the graph *G*_*i*_ and that *g* is modeled by *G*_1_ ⊛ · · · ⊛ *G*_*N*_. Let us consider what happens when enhancers 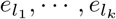 are perturbed in such a way that they are considered to no longer be working to regulate *g*. We will use a similar notation to that for the fractional change in the Introduction and denote the graph that arises from this perturbation as 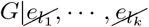. In analogy to the fractional change, we can define the *deletion effect* of the perturbation, 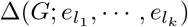, to be the proportional change in response of *g*,

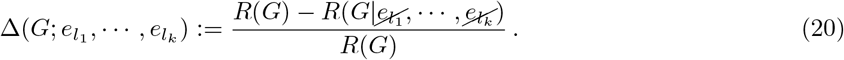

It is important to keep in mind that the deletion effect is defined in terms of a model of gene regulation, whereas the fractional change is defined in terms of experimental data. The definition in Eqn.20 implicitly assumes that the system has returned to steady state following the perturbation. Furthermore, Eqn.20 says nothing about how the enhancer is perturbed or whether a CRISPRi perturbation has the same effect as a genetic deletion.

To calculate the deletion effect using Eqn.20, we need to know the perturbed graph, 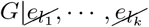. We assume that the graph shows *deletion fidelity*, which implies that the perturbation completely abrogates the function of the targeted enhancers and does not influence other enhancers. Let *m*_1_, · · ·, *m*_*p*_ be the remaining indices in 1, · · ·, *N* after *l*_1_, · · ·, *l*_*k*_ have been removed.

5. Deletion fidelity. The regulatory graph of 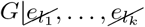 is the graph product 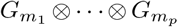, and the production rates in 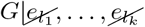 are directly inherited from *G*,

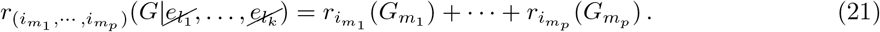

Deletion fidelity ensures that if *G* obeys Assumptions 1-4, then 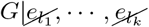 also obeys Assumptions 1-4 for the remaining enhancers 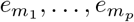 and that, for the copy-number graphs,

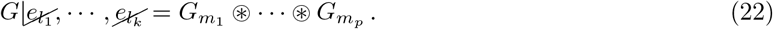

Using Eqn.22, it follows from Eqn.19 that,

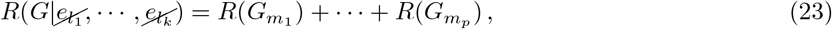

and so the formula for the deletion effect in Eqn.20 tells us that,

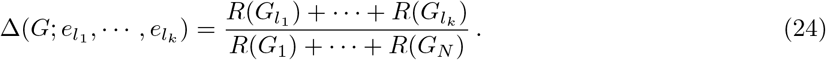

Eqns.23 and 24 are general properties that hold for the default model whenever Assumption 5 of deletion fidelity also holds. They allow us to formalise the notion of enhancer *additivity*, which we will discuss below, but, first, let us turn to the ABC Score formula.

#### The Independent-Activation-Communication (IAC) model

In the default model, the graph representing each individual enhancer can be arbitrarily complicated. To show how the ABC Score formula can arise from the default model, we need to impose the further assumption that each enhancer is modeled by the activation-communication coarse graining shown in Figs.1a and 1b.

6. The activation-communication coarse-graining. Enhancer *e*_*i*_ is described by the graph *H*_*i*_, where *H*_*i*_ is the same graph as *H* in Fig.2d. Specifically, *H*_*i*_ is the graph product of an activation graph, *K*_*a,i*_, with labels *k*_*a,i*_, *l*_*a,i*_, and a communication graph, *K*_*c,i*_, with labels *k*_*c,i*_, *l*_*c,i*_ (Fig.1c), and *H*_*i*_ = *K*_*a,i*_ ⊗ *K*_*c,i*_. The only non-zero production rate of *H*_*i*_ occurs in the state in which the enhancer is both active and communicating, where the rate is *r*_*i*_.

The overall regulatory system is then described by *G* = *H*_1_ ⊛ · · · ⊛ *H*_*N*_ (Fig. 4, Fig.S1). We call the model obeying Assumptions 1-6 the Independent-Activation-Communication (IAC) model. It follows from Eqn.16 that the response of enhancer *i* in the IAC model is given by

**Figure 4:**
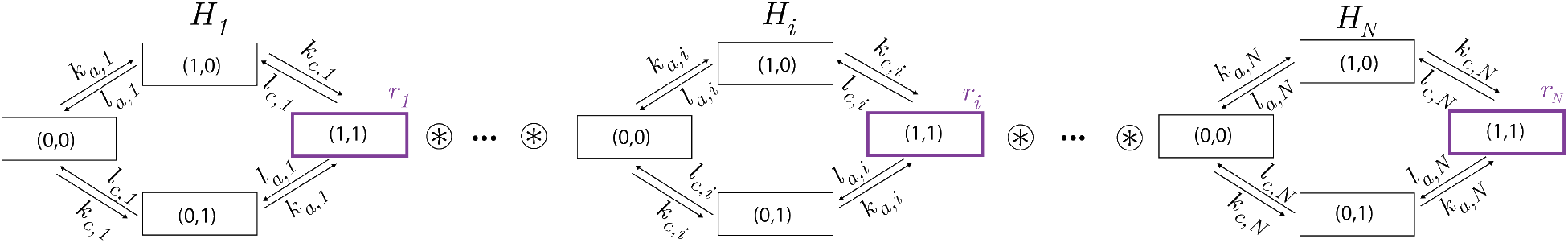
The Independent-Activation-Communication (IAC) model. A gene described by the IAC model follows Assumptions 1-6 for the component graphs *H*_1_, …, *H*_*N*_. The ordered pair of binary digits for the vertices in each *H*_*i*_ represent the activation and communication status, respectively, of each enhancer. Each *H*_*i*_ has the same structure as the graph in Fig.2d, but different labels, and represents the independence of activation and communication within each enhancer. Each *H*_*i*_ is assumed to have the same degradation rate which is omitted for clarity. See also Fig.S1.

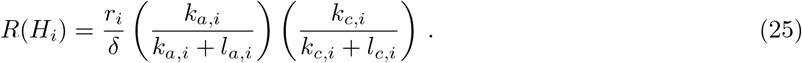

Let us define 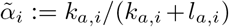 and 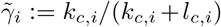 recall from Eqn.16 that these quantities are the probability of activation and the probability of communication, respectively, of enhancer *e*_*i*_. According to the fundamental property of the default model in Eqn.19, the response, *R*(*G*), of the overall graph, *G* = *H*_1_ ⊛ · · · ⊛ *H*_*N*_, is given by,

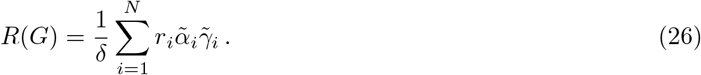

Furthermore, as a consequence of deletion fidelity (Assumption 5), it follows from Eqn.24 that the deletion effect for enhancer *e*_*q*_ is given by,

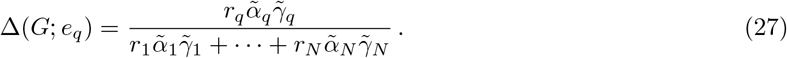

Eqn.27 shows a striking algebraic similarity to the ABC Score formula in Eqn.2. The quantity 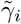, which is the probability of communication, is analogous to the ‘frequency of contact’, *γ*_*i*_, that was envisaged for the ABC model [11] and appears in Eqn.2. There are different possible interpretations for the other terms. One potential interpretation for the term 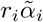in Eqn.27, which is the production rate multiplied by the probability of activation, is that it is analogous to the ‘strength of the enhancer’, *α*_*i*_, appearing in Eqn.2. Another potential interpretation is that 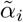 corresponds to *α*_*i*_ and that the production rates *r*_*i*_ are not represented in the ABC model. If the production rates are assumed to be equal, they would cancel out in Eqn.2, which would be consistent with a correspondence between 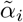and *α*_*i*_. Such interpretational ambiguities are to be expected because the ABC model is informal, while the IAC model presented here is formal. Moreover, our formal model separately specifies the regulatory state of the enhancer and its effect on transcription, whereas the ABC model does not make this distinction. An interesting question arises as to how numerical values can be assigned to the terms in Eqn.27, and how this may differ from the strategies used in [11] to give numerical values to the terms in Eqn.2, but this is an area for future work.

Eqn.27 is our clarification of the algebraic structure of the ABC Score formula. Eqn.27 rigorously follows if enhancers collectively regulate a gene according to the IAC model (Assumptions 1 to 6).

### Enhancer additivity and departures from it

The default model, satisfying Assumptions 1 to 4, exhibits *response additivity*, as shown by Eqn.19: the response of the gene to all the enhancers acting collectively is just the sum of the responses to each individual enhancer. When the default model also obeys Assumption 5 of deletion fidelity, then response additivity has a counterpart in the deletion effect, as defined in Eqn.20. This allows us to rigorously define the properties of *super-additivity* and *sub-additivity*. These departures from the properties of the default model may be helpful to interpret the effects of experimental perturbations, such as genetic deletions or CRISPRi, in which subsets of enhancers are prevented from influencing a gene and the effect of these perturbations on the gene expression response is measured.

With Assumptions 1 to 5, if *U*_1_, · · ·, *U*_*m*_ are pairwise disjoint subsets of enhancers, so that *U*_*i*_ ⊆ {*e*_1_, · · ·, *e*_*N*_} and *U*_*i*_ ∩ *U*_*j*_ = ∅ when *i* ≠ *j*, then it follows from Eqn.24 that the effect of deleting all the subsets together is just the sum of the individual deletion effects,

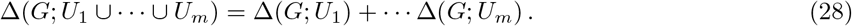

We refer to this property as *deletion additivity*. Furthermore, it is evident from Eqn.24 that, if all the enhancers are deleted, so that *U*_1_ ∪ · · · ∪ *U*_*m*_ = {*e*_1_, · · ·, *e*_*N*_}, then the *total deletion effect* must be 1,

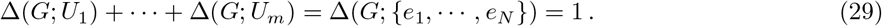

Assuming deletion fidelity, the total deletion effect being 1 is equivalent to the response additivity in Eqn.19. A special case of Eqn.29 arises if all enhancers are deleted individually, when, once again, the total deletion effect is 1,

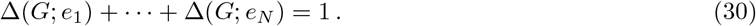

Now suppose that a gene *g* is regulated by *N* enhancers, *e*_1_, · · ·, *e*_*N*_, each enhancer is modeled by the graph *G*_*i*_ and the regulatory graph of *g, G*, has the product structure, *G*_1_ *×* · · · *× G*_*N*_. The labels in *G* need not be related to those of the component graphs *G*_*i*_, so that *G* need not be the product graph *G*_1_ ⊗· · · ⊗ *G*_*N*_. We can no longer calculate *R*(*G*) in terms of *R*(*G*_*i*_). However, we can still define through Eqn.20 the deletion effect ∆(*G*; *U*) for any collection *U* ⊆ {*e*_1_, · · ·, *e*_*N*_} of enhancers. We say that *g* exhibits response *super-additivity* if,

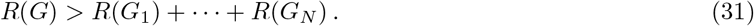

In terms of the deletion effect, this corresponds to when a collective deletion has less effect than the sum of the individual deletions, so that,

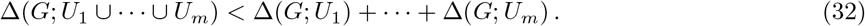

Similarly, *g* exhibits response *sub-additivity* if,

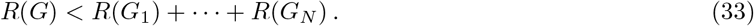

and this corresponds to the collective deletion having more effect that the sum of the individual deletions,

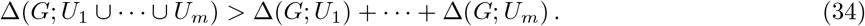

Experimentally, response additivity [15, 23, 38–43], super-additivity [15, 21, 39–44] and sub-additivity [38, 40, 41] have all been observed. Because the super-additive and sub-additive findings cannot be accounted for by the default model with deletion fidelity, we next consider some extensions of this model that show how such effects could arise.

### Mechanisms beyond the default model

In the following sections, we examine two departures from the default model and consider their impact on whether enhancers act additively (Eqn.19), super-additively (Eqn.31) or sub-additively (Eqn.33). This will also illustrate how our modeling framework can be used to reason about different biological mechanisms.

#### Non-additivity in mRNA production rates

In the default model, the summation of production rates in Assumption 3 is crucial for the property of enhancer additivity in Eqn.19. The production rate is a convenient abstraction that aggregates over many underlying molecular mechanisms, such as RNA Polymerase recruitment, pausing and elongation. It is conceivable that, when multiple enhancers jointly influence transcription, the resulting rate is a more complex function than simple addition [38]. Here, we consider the effect of dropping Assumption 3.

Let us assume that we have two enhancers, *e*_1_ and *e*_2_, which are described by the graphs *H*_1_ = *K*_*a*,1_ ⊗*K*_*c*,1_ and *H*_2_ = *K*_*a*,2_ ⊗ *K*_*c*,2_, respectively, as specified in Assumption 6 in the coarse-grained version of our default model. The overall regulatory graph is given by *H*_1_ ⊗ *H*_2_, so that *e*_1_ and *e*_2_ remain independent (Assumption 2). Note that (*K*_*a*,1_ ⊗ *K*_*c*,1_) ⊗ (*K*_*a*,2_ ⊗ *K*_*c*,2_) has a product hierarchy and its vertices are therefore indexed by tuples of tuples of the form,

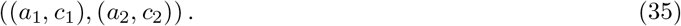

Here, *a*_*i*_ and *c*_*i*_, for *i* = 1, 2, are coordinates for activation and communication, respectively, which take the values 0 and 1 in all cases. The graphs *H*_1_ and *H*_2_ have mRNA production rates, *r*_1_ and *r*_2_, respectively, as specified in Eqn.15 and mRNA degradation rate *δ*.

We now consider a copy-number graph, *G*^⋄^, whose regulatory graph is given by (*K*_*a*,1_ ⊗*K*_*c*,1_)⊗(*K*_*a*,2_ ⊗*K*_*c*,2_) but whose production rates do not obey Assumption 3. Note that we use the same symbol, *G*^⋄^, for the regulatory graph and the copy-number graph and rely on the context to clarify which is meant. There are many ways to assign production rates to the vertices of *G*^⋄^ which do not obey Assumption 3; here we consider one of the simplest possible ways. We define 3 subsets of vertices of *G*^⋄^ in terms of the coordinates in Eqn.35: *W* := {*a*_1_ = 1, *c*_1_ = 1, *a*_2_ = 1, *c*_2_ = 1}, *U* := {*a*_1_ = 1, *c*_1_ = 1} *\ W* and *V* := {*a*_2_ = 1, *c*_2_ = 1} *\ W*. We assign the production rate of vertices in *U* to be *r*_1_, of vertices in *V* to be *r*_2_ and of the vertex in *W* to be (1 + *μ*)(*r*_1_ + *r*_2_); all other vertices have production rate 0. We can summarise these assumptions in the following table, which gives the production rate for each of the 16 states in *G*^⋄^ in the coordinate system described by Eqn.35.

| state | rate | state | rate |
| --- | --- | --- | --- |
| $((0, 0), (0, 0))$ | 0 | $((1, 0), (0, 0))$ | 0 |
| $((0, 0), (0, 1))$ | 0 | $((1, 0), (0, 1))$ | 0 |
| $((0, 0), (1, 0))$ | 0 | $((1, 0), (1, 0))$ | 0 |
| $((0, 0), (1, 1))$ | $r_2$ | $((1, 0), (1, 1))$ | $r_2$ |
| $((0, 1), (0, 0))$ | 0 | $((1, 1), (0, 0))$ | $r_1$ |
| $((0, 1), (0, 1))$ | 0 | $((1, 1), (0, 1))$ | $r_1$ |
| $((0, 1), (1, 0))$ | 0 | $((1, 1), (1, 0))$ | $r_1$ |
| $((0, 1), (1, 1))$ | $r_2$ | $((1, 1), (1, 1))$ | $(1 + \mu)(r_1 + r_2)$ |

We further assume that *μ ≥* −1 to ensure that the production rate of *W* does not become negative. If *μ* = 0, then Assumption 3 holds for *G*^⋄^ but not otherwise. We also assume that *G*^⋄^ has degradation rate *δ*. We now calculate *R*(*G*^⋄^). Using the Sanchez-Kondev theorem in Eqn.14 we have,

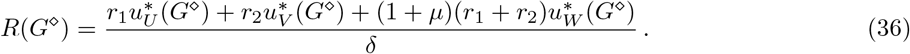

Expanding the term on the right hand side of Eqn.36 and rearranging terms results in,

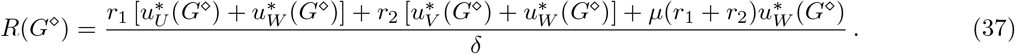

Using the fact that the pairs of sets *U* and *W*, and, *V* and *W* are disjoint gives,

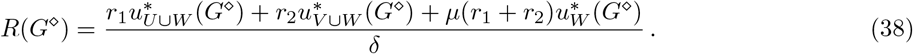

We now note that, by definition, *U* ∪*W* = {*a*_1_ = 1, *c*_1_ = 1} and *V* ∪*W* = {*a*_2_ = 1, *c*_2_ = 1}. Given the independence assumption on *G*^⋄^, we can apply Eqn.11 and have that 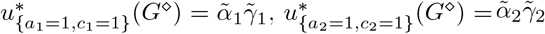 and 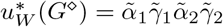. Substituting into Eqn.38, we have,

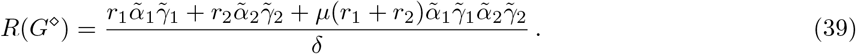

Given that

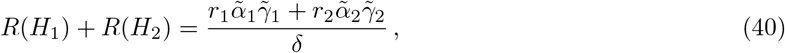

we see that *e*_1_ and *e*_2_ act additively for *μ* = 0 (Eqn.19), act super-additively for *μ >* 0 (Eqn.31) and act sub-additively for *μ* ∈ [−1, 0) (Eqn.33).

#### Non-independence in regulatory transitions between enhancers

So far all the graphs we have considered obey regulatory independence as defined by Assumption 2: the state of one enhancer does not affect the transitions or the rates of any other enhancer. Let us examine this more closely for the IAC model, with just two enhancers, *e*_1_ and *e*_2_, described by graphs *H*_1_ and *H*_2_, respectively, as in the previous subsection. According to Assumption 6, *H*_1_ = *K*_*a*,1_ ⊗ *K*_*c*,1_ and *H*_2_ = *K*_*a*,2_ ⊗ *K*_*c*,2_ and the overall regulatory system is therefore described by the graph, *G* = *H*_1_ ⊗ *H*_2_ = (*K*_*a*,1_ ⊗ *K*_*c*,1_) ⊗ (*K*_*a*,2_ ⊗ *K*_*c*,2_). Using Eqn.12 and the coordinate system in Eqn.35, we have that 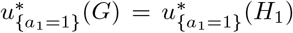 and 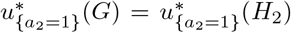. That is, the probability of activation of an enhancer does not depend on the presence of the other enhancer. However, if probability of activation is measured by H3K27ac ChIP-Seq, there is evidence that perturbation of a single enhancer can result in altered H3K27ac signal at distal enhancers [15, 22, 45, 46]. If such changes at the distal enhancer are not caused by the perturbation method itself, so that the perturbation obeys the fidelity conditions in Assumption 5, then such experiments suggest that there may be non-independence between enhancers at the level of activation.

In order to model non-independence between enhancers, we will consider a gene to be modeled by the graph *G*^*♯*^, where *G*^*♯*^ is the product between an activation graph *A*^*♯*^ and a communication graph *C*, so that *G*^*♯*^ = *A*^*♯*^ ⊗ *C* (Fig.5). *A*^*♯*^ has the structure *K*_*a*,1_ × *K*_*a*,2_, in which *K*_*a*,1_ and *K*_*a*,2_ are both present as the subgraphs,

**Figure 5:**
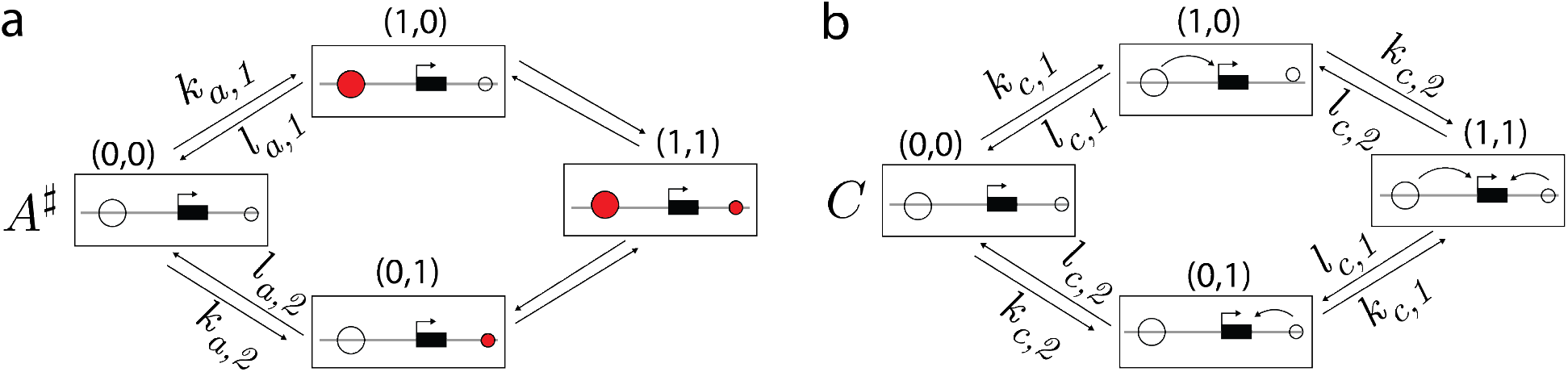
A model of non-independence in activation between enhancers. **(a)** The graph *A*^*♯*^ represents the activation components of each enhancer. *A*^*♯*^ has the structure of *K*_*a*,1_ *× K*_*a*,2_. Labels on the unmarked edges can be arbitrary. **(b)** The graph *C* = *K*_*c*,1_ ⊗ *K*_*c*,2_ represents the communication components of each enhancer.

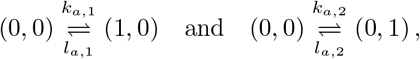

respectively (Fig.5a). The remaining labels, on the edges (1, 0) ⇌ (1, 1), which specify the rates of activation and deactivation of *e*_2_ when *e*_1_ is activated, and on the edges (0, 1) ⇌ (1, 1), which specify the rates of activation and deactivation of *e*_1_ when *e*_2_ is activated, can be arbitrary. For simplicity, we assume that *C* = *K*_*c*,1_ ⊗ *K*_*c*,2_, (Fig.5b), but note that non-independence in communication could be considered similarly. *G*^*♯*^ has the same structure as *H*_1_ × *H*_2_, and thus still models the activation and communication statuses of *e*_1_ and *e*_2_, but using the form *G*^*♯*^ = *A*^*♯*^ ⊗ *C* allows us to clarify the independence relationships in *G*^*♯*^. Under this reorganization, the vertex in Eqn.35 is now described in a new coordinate system as,

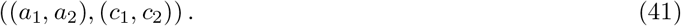

We assume that *G*^*♯*^ has the same mRNA production rates as for the IAC model. In terms of the vertex subsets *W* = {*a*_1_ = 1, *c*_1_ = 1, *a*_2_ = 1, *c*_2_ = 1}, *U* = {*a*_1_ = 1, *c*_1_ = 1} *\ W* and *V* = {*a*_2_ = 1, *c*_2_ = 1} *\ W*, the vertices in *U* have production rate *r*_1_, those in *V* have production *r*_2_ and those in *W* have production rate *r*_1_ + *r*_2_; all other vertices have production rate 0. We can summarise this in the following table, in which the states are described by the coordinate system in Eqn.41.

| state | rate | state | rate |
| --- | --- | --- | --- |
| $((0, 0), (0, 0))$ | 0 | $((1, 0), (0, 0))$ | 0 |
| $((0, 0), (0, 1))$ | 0 | $((1, 0), (0, 1))$ | 0 |
| $((0, 0), (1, 0))$ | 0 | $((1, 0), (1, 0))$ | $r_1$ |
| $((0, 0), (1, 1))$ | 0 | $((1, 0), (1, 1))$ | $r_1$ |
| $((0, 1), (0, 0))$ | 0 | $((1, 1), (0, 0))$ | 0 |
| $((0, 1), (0, 1))$ | $r_2$ | $((1, 1), (0, 1))$ | $r_2$ |
| $((0, 1), (1, 0))$ | 0 | $((1, 1), (1, 0))$ | $r_1$ |
| $((0, 1), (1, 1))$ | $r_2$ | $((1, 1), (1, 1))$ | $r_1 + r_2$ |

As in the previous section, we can use the Sanchez-Kondev theorem in Eqn.14 to calculate,

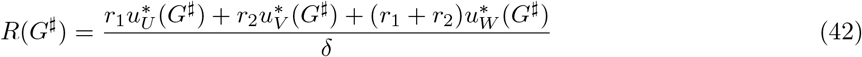

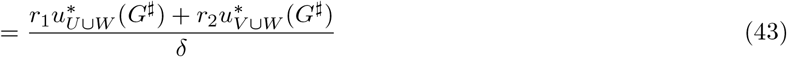

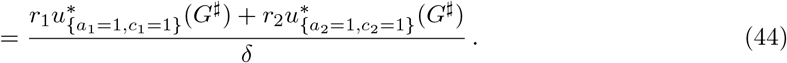

Given that *G*^*♯*^ = *A*^*♯*^ ⊗ *C*, the probabilities of the sets {*a*_1_ = 1, *c*_1_ = 1} and {*a*_2_ = 1, *c*_2_ = 1} factor according to Eqn.11. Continuing from Eqn.44 we have,

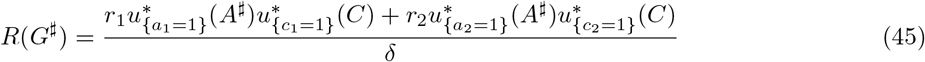

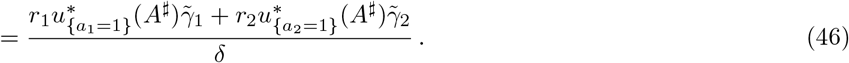

Eqn.46 shows that the labels of *A*^*♯*^ do not directly appear in *R*(*G*^*♯*^); they only affect *R*(*G*^*♯*^) through the enhancer activation probabilities 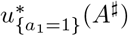 and 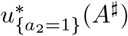. As in the previous section, we note that the sum of the individual enhancer responses is,

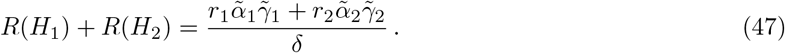

Comparing Eqns.46 and 47, we see that whether the enhancers act additively (Eqn.19), sub-additively (Eqn.33) or super-additively (Eqn.31) depends on the terms

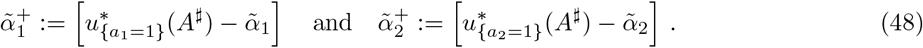

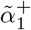 represents the change in the probability of activation of enhancer 1 due to the presence of enhancer 2, and 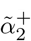 represents the change in the probability of activation of enhancer 2 due to the presence of enhancer 1. If both 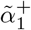 and 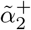 are positive, then *e*_1_ and *e*_2_ act super-additively; if they are both negative, then *e*_1_ and *e*_2_ act sub-additively. If 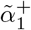 and 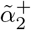 are of different signs, then the enhancers may act super-additively or subadditively depending on the relative magnitude of these terms compared to 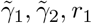 and *r*_2_. Experimental data in which both 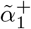 and 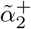 have been measured is limited. There are experimentally observed instances in which both of these terms are positive [15, 45] but the precise form that the graph *A*^*♯*^ takes in these cases is unknown. We are unaware of experiments that have observed 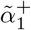 and 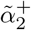 of differing signs. Whether the experimentally observed non-additivity between enhancers can be explained by the non-independence between enhancers as described in this section is a future area of research.

## Discussion

In this paper, we have introduced mathematical formulations of a *default model* and an *IndependentActivation-Communication model* (IAC model) for how multiple enhancers collectively regulate a gene. The default model encodes the notion that enhancers operate independently of each other (Assumptions 2 and 3). At the same time, the default model imposes no assumptions on how the individual enhancers themselves are working, at the level of transcription factors, co-regulators, chromatin, etc. They can be arbitrarily complicated, so long as they operate within the Markovian setting that is commonly assumed for analysing gene regulation. The default model assumptions imply that the collective response of a gene, as measured by the mean mRNA level, is the sum of the responses coming from each enhancer individually, which we have called *response additivity* (Eqn.19). The default model explains the mechanistic requirements for a gene to exhibit this property and clarifies how ‘independence implies additivity’. We emphasize that independence refers here to assumptions about gene regulatory mechanisms whereas additivity refers to the consequences of those assumptions on steady-state gene expression. The default model supports the view that response additivity is a reasonable baseline against which to assess the collective action of enhancers in regulating a gene.

One of the advantages of the default model is that, because it is mathematically formulated, it allows mechanistic departures from its assumptions to be systematically analysed. We have shown how departures from Assumptions 2 and 3 can give rise to response super-additivity (Eqn.32) as well as sub-additivity (Eqn.34). As we have noted, response additivity [15, 23, 38–43], super-additivity [15, 21, 39–44] and subadditivity [38, 40, 41] have all been observed experimentally. The default model suggests the mechanistic assumptions that could be experimentally tested to determine what underlies the observed response behaviour.

The IAC model is a special case of the default model that further assumes deletion fidelity, which allows enhancers to be removed from the collective without influencing the remaining enhancers (Assumption 5), and also assumes that individual enhancers can be described at a coarse-grained level in which they are independently becoming activated and communicating their state to the gene (Assumption 6). Under Assumptions 1 to 6 for the IAC model, we derive a formula for the deletion effect of an individual enhancer (Eqn.27) that shows a striking algebraic relationship to the ABC Score formula in Eqn.2. This relationship suggests that the IAC model has accurately captured in mathematical terms the core intuitions behind the ABC model from which the Score formula emerged [11].

A persistent conceptual theme that underlies the results reported here is that of independence. The default model assumes that enhancers act independently, both in their regulatory state (Assumption 2) and in their effect on mRNA production (Assumption 3). Furthermore the IAC model assumes that individual enhancers become activated and communicating independently (Assumption 6). Our clarification of the

ABC Score formula thus arises from assuming independence *between* enhancers, along with independence of activation and communication *within* each enhancer. The product construction on graphs and on graph structures has been the key mathematical tool for rigorously defining independence, illustrating the value of the graph-based linear framework for analysing gene regulation. We note that the concept of deletion fidelity (Assumption 5) is also easily defined in the context of graphs.

Previous work used finite linear framework graphs to describe gene regulation [26]. Here, we have introduced *copy-number graphs*, which have infinitely many vertices that keep track of both regulatory states as well as the numbers of expressed mRNAs (Fig.2). Copy-number graphs allowed us to exploit the Sanchez-Kondev theorem and calculate the mean mRNA number at steady state in terms solely of the finite regulatory graph (Eqn.14). We therefore avoided dealing with infinite graphs despite relying on them. Importantly, the linear framework also allows the unknown parameters within graphs to be treated symbolically, so that conclusions may be drawn, as we saw above, without the need for assigning numerical values to any of the parameters.

Beyond its utility in clarifying the ABC Score formula, the activation-communication coarse graining in Assumption 6 provides an interesting lens through which to investigate enhancers. Many new experimental technologies have emerged which allow perturbing entire enhancer sequences as a whole (as opposed to small changes to DNA within an enhancer sequence). Such technologies include synthesizing and integrating long DNA sequences [43, 44], modulating the genomic position of an enhancer [47–51], high throughput enhancer perturbations with CRISPRi [11–13] and combining CRISPRi screens with rapid protein degradation [52]. By considering which perturbations affect, and do not affect, activation and communication, it may be possible to probe the validity of the activation-communication coarse graining itself.

As noted in the Introduction, the ABC Score formula has been widely adopted for predicting enhancer-gene connections. It has also been suggested that it could be combined with other predictive methods [53] and that the ABC model could be used as a guiding principle in formulating other quantitative models [54]. We believe the mathematical formulations that we have introduced here provide a foundation for such efforts.

The ABC Score formula is quantitative (Eq.2) but the ABC model that gave rise to it is not a formal mathematical model but, rather, an informal statement about the features, of enhancer activation and contact, that are believed to be important in determining the response of a gene. Such informal models play a critical role in biology but have the disadvantage that the underlying mechanistic requirements are not clear. It is therefore difficult to know when the model can be applied and what can be deduced from it when it does apply. In contrast, the mechanistic assumptions underlying our formal mathematical models are precisely stated—Assumptions 1 to 4 for the default model and Assumptions 1 to 6 for the IAC model— making it clear when the model can be applied and suggesting experimental tests to check the assumptions. Moreover, if those assumptions are met, then the conclusions we have drawn, such as the response additivity of the default model (Eqn.19) and the enhancer deletion formula for the IAC model (Eqn.27), are guaranteed to hold as a matter of mathematical logic [55]. If those conclusions are not found experimentally, for example, if response additivity is not found, then we know, as a matter of logic, that at least one of the assumptions underlying the corresponding model does not hold. This understanding can, in turn, inform experiments to determine where the departures from the assumptions occur. Such an approach allows a level of rigorous reasoning about enhancer behaviour in gene regulation that is significantly harder to undertake with only an informal quantitative model.

Mathematical theory has typically been introduced to analyse data, but the conceptual issues underlying gene regulation are sufficiently intricate that theory may be necessary to understand the kinds of experiments that are needed and how the data from them should best be interpreted [56]. Studies of the simple repression motif in bacterial gene regulation may have already reached that point [57–60], as reviewed in [61]. The foundation provided here, based on the linear framework, may offer similar opportunities in the eukaryotic context. We believe our rigorous mathematical approach can play a significant role in investigating the intricate interplay of enhancers in regulating gene expression.

## Methods

### A graph theory interpretation of the Sanchez and Kondev theorem

In this section we provide a proof of Eqn.14. We follow the Sanchez and Kondev approach described in but present it using the graph theory notation and language used in this paper. Sanchez and Kondev provide in [34] a recurrence relation for all the moments of the mRNA probability distribution. A graph theory interpretation of these results, together with generalisations, will be presented in a separate paper; here we focus on the first moment only. We use bold face to denote matrices and vectors.

#### The steady-state distribution of a finite graph

Let *G* be a finite regulatory graph on the vertex set *V* (*G*) = {1, …, *N*}. We define the *Laplacian matrix* of *G*, ℒ = ℒ (*G*), to be the *N × N* matrix,

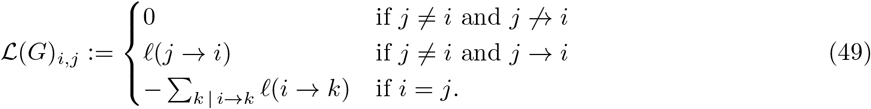

As mentioned in the main text, *G* is equivalent to a continuous-time Markov process on the state space {1, …, *N*} [25, 28, 32]. Let *u*_*i*_(*t*) be the probability that the process occupies state *i* at time *t*. Then the time evolution of the probability vector,

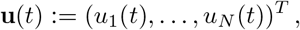

is given by the master equation

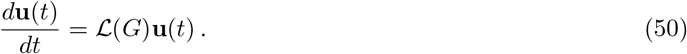

If *G* is strongly connected, then the kernel of ℒ(*G*) is one dimensional, so there is a unique vector, **u**^∗^(*G*), such that ℒ(*G*)**u**^∗^(*G*) = 0 and *u*_1_(*G*) + · · · + *u*_*N*_ (*G*) = 1. **u**^∗^(*G*) is the steady-state probability distribution on *G*.

#### The master equation for a copy-number graph

Let *G* ⋉ *P* denote a copy-number graph with regulatory graph *G*, production rate vector **r** ∈ ℝ^*N*^ and degradation rate *δ*. Let **Π** be the diagonal matrix of production rates, **Π**_*i,i*_ = *r*_*i*_ and **Π**_*i,j*_ = 0 when *i ≠j*, and let **I** be the *N × N* identity matrix. Let

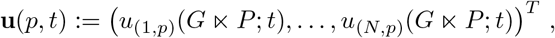

be the vector of probabilities over the regulatory states with mRNA copy number *p*. It follows from the definition of the copy-number graph in the main text that **u**(*p, t*) satisfies the master equation,

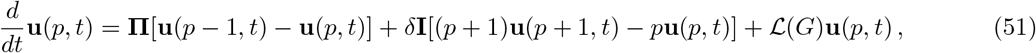

in which terms with arguments of *p* − 1 are appropriately omitted when *p* = 0. The first term of Eqn.51 arises from mRNA production, the second term from mRNA degradation and the third term from transitions in the regulatory graph. Eqn.51 can be rewritten as,

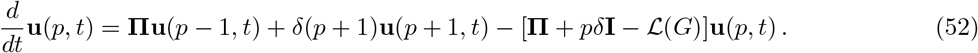

We now let

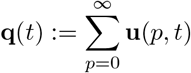

be the vector of marginal probabilities for the regulatory states. Proceeding from Eqn.52 we have,

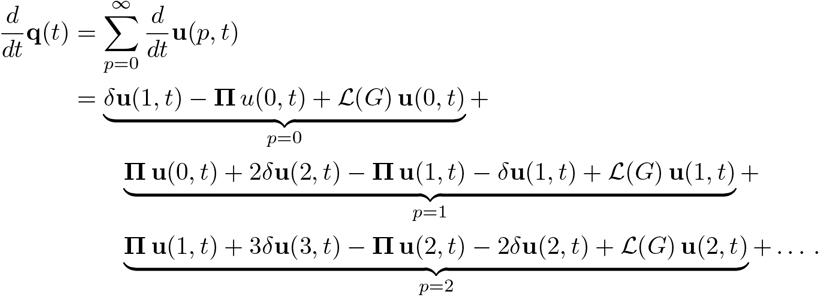

This is a telescoping sum which simplifies to

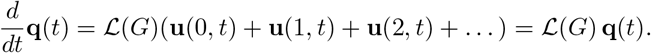

It follows that the steady-state marginal probability vector, **q**^∗^, lies in the kernel of ℒ(*G*) and must therefore be equal to **u**^∗^(*G*),

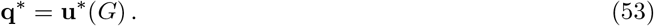

In other words, the steady-state marginal distribution of regulatory states in a copy-number graph is identical to the steady-state distribution of regulatory states in a finite regulatory graph.

**Proof of Eqn.14**

We want to show that,

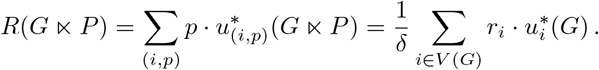

Let **u**^∗^(*p*) denote the steady-state probability distribution over the copy-number graph. (Note the distinction with the marginal probability distribution over the regulatory states, *u*^∗^(*G*) = ∑_*p*_ *u*^∗^(*p*).) Let

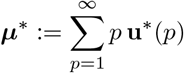

be the corresponding steady-state average copy number vector. Evidently,

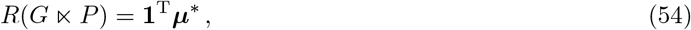

where **1** is the all-ones column vector of dimension *N*. Now let

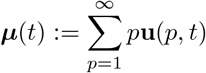

be the time-dependent average copy-number vector. It follows from Eqn.52 that,

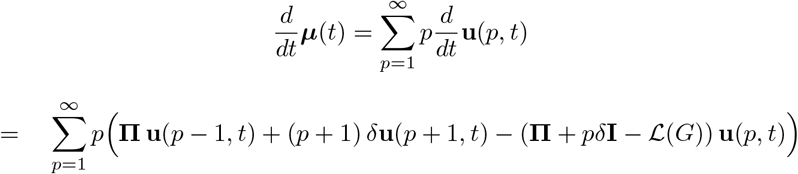

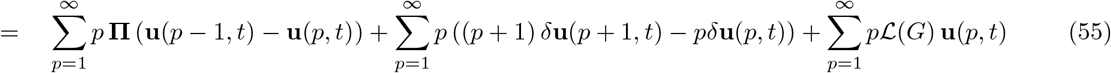

The first summand in Eqn.55 can be simplified to,

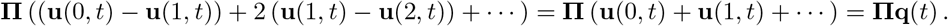

The second summand can be simplified to,

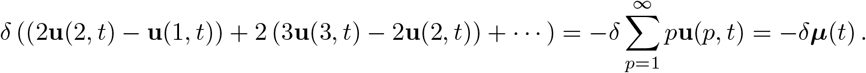

And the third summand is evidently just *ℒ* (*G*) ***μ***(*t*). Combining these three simplifications, we see that,

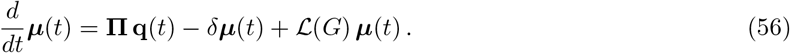

At steady state this becomes,

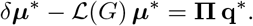

Multiplying both sides **1**^T^, and recalling that,

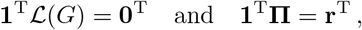

we find that,

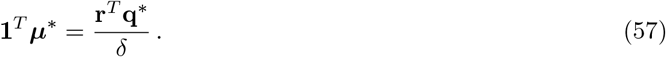

Using Eqns.53 and 54, we see that Eqn.57 becomes,

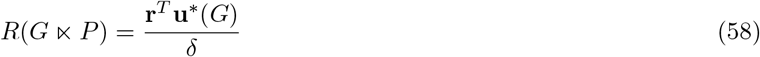

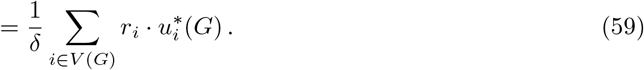

as required. This completes the proof of Eqn.14.

### Proof of response summation in the default model

In this section we prove Eqn.19 which shows that, within the default model, the collective response of all the enhancers is the sum of their individual responses. That is, if *G* = *G*_1_ ⊛ · · · ⊛ *G*_*N*_, then

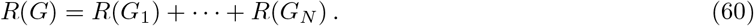

We consider the case with only two enhancers, *N* = 2, from which the general case follows easily. Recall from the Sanchez and Kondev formula in Eqn.14 that

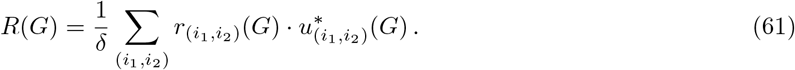

Assumption 3 on response summation tells us that 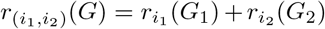 and Eqn.8 for the product graph tells us that 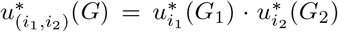. Substituting these expressions into Eqn.61 gives the following formula for *δ* · *R*(*G*),

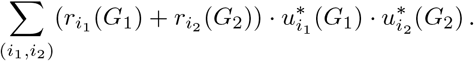

We can perform the summation over (*i*_1_, *i*_2_) in any order, for instance by first summing over *i*_2_ and then summing over *i*_1_. This gives,

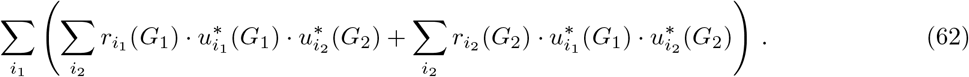

In the inner left-hand sum over *i*_2_, the terms indexed by *i*_1_ are constant and may be extracted from that sum to give

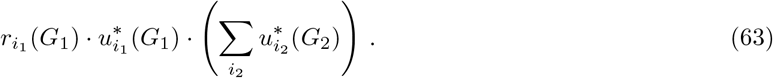

Total probability always sums to 1, so that 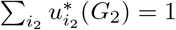. and Eqn.63 reduces to

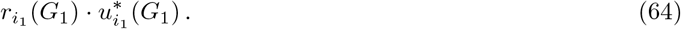

Similarly, the inner right-hand sum in Eqn.62 may be written as

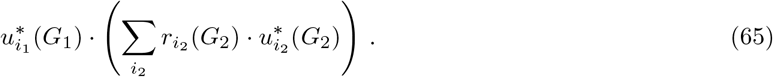

We recognise from Eqn.14 that the sum in brackets is the response of graph *G*_2_, so that Eqn.65 becomes,

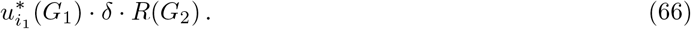

We can now substitute Eqns.64 and 66 back into Eqn.62 to get,

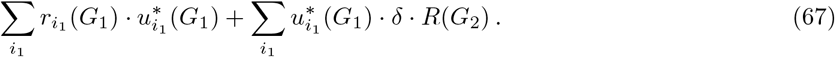

We recognise from Eqn.14 that the left-hand sum is *δ* times the response of graph *G*_1_. In the right-hand sum, we can extract the terms that do not depend on *i*_1_ and use once again that the total probability is 1. This allows us to rewrite Eqn.67 as,

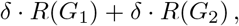

from which we conclude that, indeed,

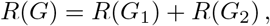

as claimed. This completes the proof of Eqn.19.

### Symbolic computations

We have provided mathematical proofs for all of our results. However, many of our results were originally discovered by exploration using computer algebra systems. Specifically, the Sage [62] computer algebra system, and the SymPy [63] and NetworkX [64] Python packages were crucial for the development of this paper.

## Acknowledgements

JN, K-MN and JG were funded in part by NIH award R01GM122928; JN was also funded the NSF-Simons Center for Mathematical and Statistical Analysis of Biology at Harvard University. We thank Zeba Wunderlich, Rosa Martinez-Corral, Jané Kondev and members of the Gunawardena lab for discussions and comments on the manuscript.

---

**Figure S1:**
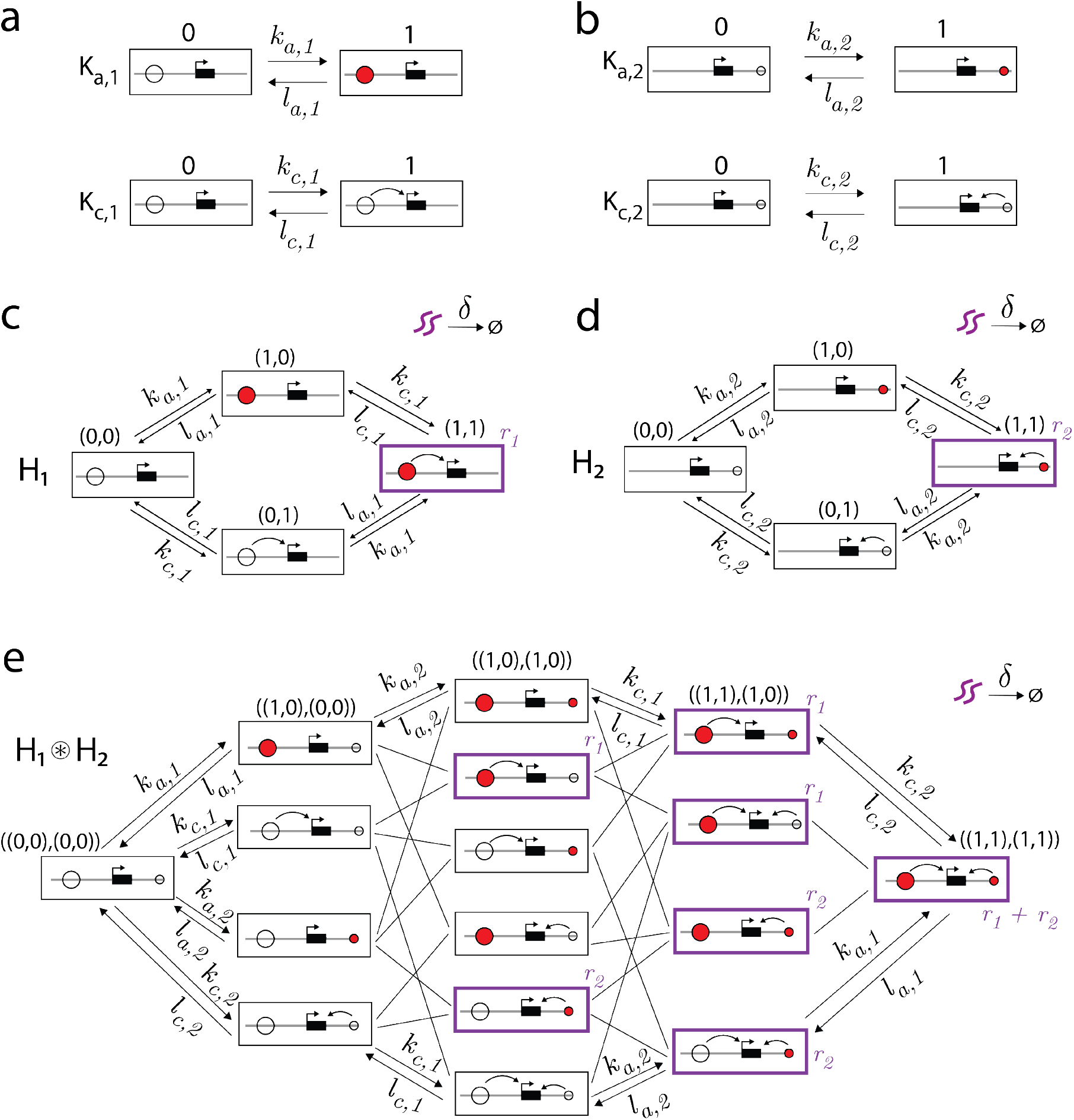
IAC model for *N* = 2 enhancers as a product graph of product graphs. **(a)** Graphs describing the activation and communication status of enhancer 1. **(b)** Graphs describing the activation and communication status of enhancer 2. **(c)** The graph *H*_1_ whose underlying regulatory graph is *K*_*a*,1_ ⊗ *K*_*c*,1_. **(d)** The graph *H*_2_ whose underlying regulatory graph is *K*_*a*,2_ ⊗ *K*_*c*,2_. **(e)** The graph *H*_1_ ⊛ *H*_2_ satisfying the default model Assumptions 1-4 with components *H*_1_ and *H*_2_. The regulatory graph of *H*_1_ ⊛ *H*_2_ is given by (*K*_*a*,1_ ⊗ *K*_*c*,1_) ⊗ (*K*_*a*,2_ ⊗ *K*_*c*,2_). Each vertex of this graph corresponds to the activation and communication statuses of both enhancers. For the vertices along the top of the graph, the product-graph binary notation is also provided using the coordinate system ((*a*_1_, *c*_1_), (*a*_2_, *c*_2_)). Reverse edges and labels of most edges are omitted for clarity. Production states are highlighted in purple with corresponding production rates also in purple font.

## Notes

### Competing Interest Statement

The authors have declared no competing interest.

